# Adaptation, Specialization, and Coevolution within Phytobiomes

**DOI:** 10.1101/120493

**Authors:** David A. Baltrus

**Affiliations:** School of Plant Sciences, University of Arizona, Tucson AZ 85721; School of Animal and Comparative Biomedical Sciences, University of Arizona, Tucson AZ 85721

## Abstract

Growth patterns of individual plants and evolutionary trajectories of plant communities are intimately linked with and are critically affected by host-associated microbiomes. Research across systems has begun to shed light on how these phytobiomes are established and under laboratory and natural conditions, and have cultivated hope that a better understanding of the governing principles for host-microbe interactions can guide attempts to engineer microbiomes to boost agricultural yields. One important, yet relatively understudied, parameter in regards to phytobiome membership is the degree to which specialization and coevolution between plant species and microbial strains structures these communities. In this article, I provide an broad overview about current knowledge concerning mechanisms enabling adaptation and specialization of phytobiome communties to host plants as well as the potential for plants themselves to recruit and cultivate interactions with beneficial microbes. I further explore the possibility of host-beneficial microbe coevolution and suggest interactions that could promote the evolution of such close-knit partnerships. It is my hope that this overview will encourage future experiments that can begin to fill in this black box of ecological and evolutionary interactions across phytobiomes.

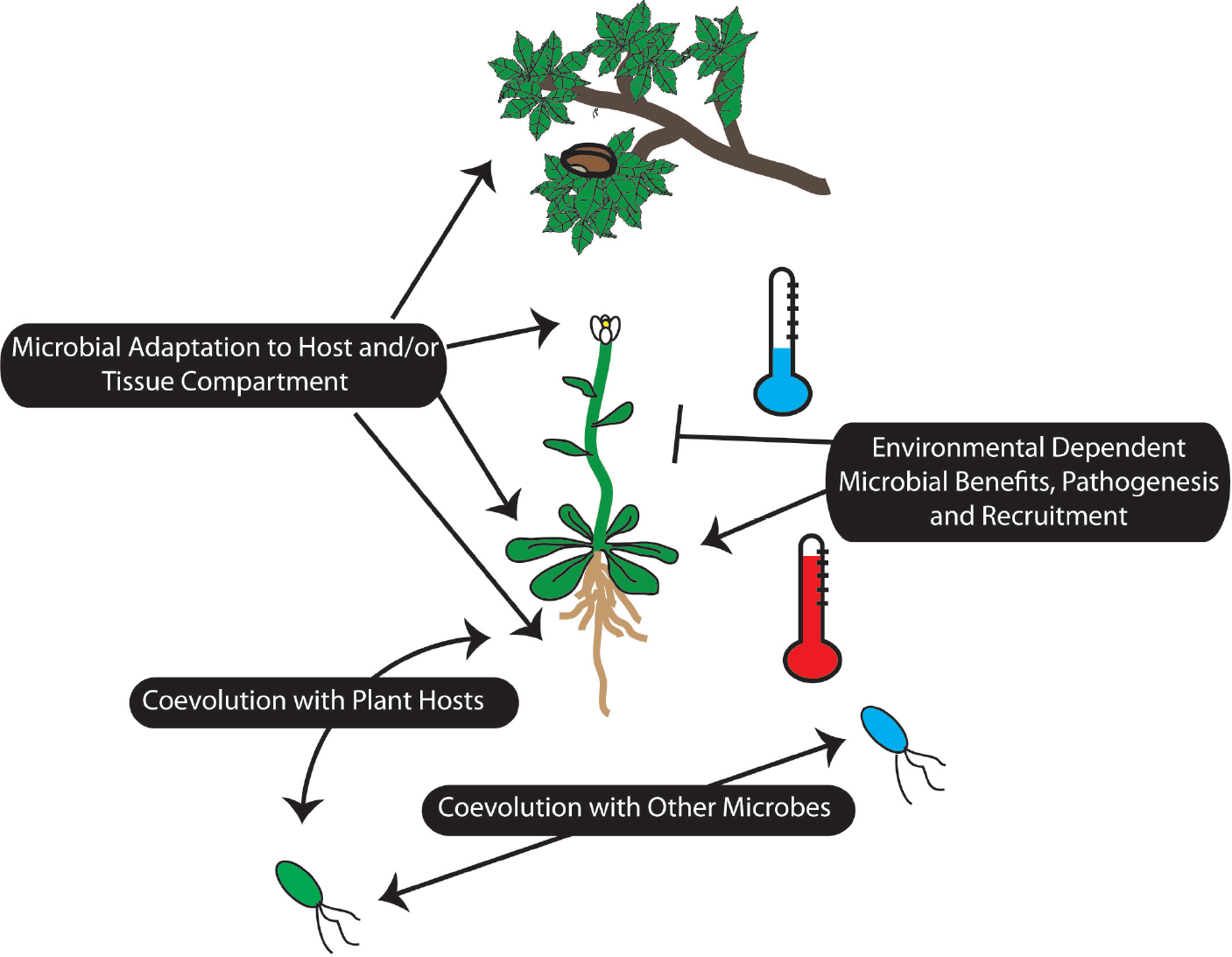

## Introduction

Recent years have seen an explosion of research focused on characterizing plant associated microbiomes, referred to collectively as the phytobiome[1–4]. This research has furthered our understanding that microbes can greatly influence host development [5–7]and bias evolutionary dynamics of plant populations[8]. Overall, promises of a new green revolution abound as ecologists race to understand factors that shape phytobiome communities and industrial researchers search for ways to efficiently engineer microbiomes to boost production[9]. Despite understandably lofty expectations and excitement for all areas of microbiome research, it is often good practice to step back to assess blind spots in research and to examine where future challenges might lie. The goal for this opinion is to begin to decorate the context of phytobiome research with a greater emphasis on evolutionary dynamics involving bacteria and plant hosts by placing a critical eye on adaptation across phytobiomes. While other microbes are part of the phytobiome, this article focuses solely on bacterial interactions for the sake of simplicity. Going forward, incorporation of evolutionary influences and limitations could help to structure ecological questions about phytobiomes while also generating new hypotheses to explain patterns of succession. Likewise, such thinking could also aid agricultural research begin to match the hype by highlighing critical bottleneck points that can be exploited to facilitate shifts in community structure.

## A Note About Phytobiome Membership

A first step for evaluating any scientific field is to find the correct context to place and evaluate new research results. One easily overlooked, but nonetheless intrinsically important, aspect to studying plant microbiomes is the realization that pathogenic strains are themselves part of all microbiomes [10] and that of the same evolutionary questions [11]likely apply to pathogens and symbionts alike. While pathogens differentiate themselves from the rest of the phytobiome by transiently blooming to large population sizes and causing tissue destruction, otherwise they may just blend into the background. This is actually a core aspect of epidemiology, as the “Disease triangle” suggests that many pathogens only cause disease under the “right” conditions[12]. Regardless of how membership within the phytobiome is classified, there exists a vast wealth of knowledge concerning molecular mechanisms and evolutionary influences of phytopathogens on host plants and this body of work should not be forgotten when discussing microbiomes writ large.

## Are Microbiomes Specialized to Particular Host Plants?

Broad surveys have demonstrated that bacteria are found in abundance across all plant tissues under natural conditions and can be acquired in a variety of ways (Box 1). While most studies are limited by sampling a small number of loci (and usually *just* 16S rDNA), one general trend that emerges is that community distributions are broadly similar across all plant compartments as three higher level taxa (Proteobacteria, Actinobacteria, and Bacteroides) dominate[13–19]. Dialing up the taxonomic resolution demonstrates that each tissue compartment does harbor its own particular mix of strains and species despite this overall stable background although phenotypic functions may be similar[16]. For example, while Pseudomonads can be found across all types of plant tissues the particular representative *Pseudomonas* OTUs (Operational taxonomic units) found in leaves will often differ from those found within roots.

### Box 1

Evolutionary interactions will be driven by the strength of associations between phytobiomes and their plant hosts. It is thus critical to explore how these communities are acquired, and to identify key parts of the life cycle when membership can shift. To this point, it may be more fruitful to focus on seed inoculation because early stages of the lifecycle could have profound downstream effects on ecological succession. Furthermore, whether microbes are vertically inherited or horizontally acquired will have profound implications for whether strains can adapt to or coevolve with specific plant hosts. A better understanding of microbiome transmission and recruitment is key if the goal is to engineer phytobiome communities, as a recent report suggests that environmentally recruited locally adapted microbes can overwhelm engineered microbiomes inoculated onto plants if given enough time.

One of the greatest challenges when discussing host-associated microbiomes is identifying specific instances where bacteria have adapted to particular host plant lineages. Definitive observations about specialization remain challenging, however, because while host range makes intuitive sense in a qualitative way real differences are often quantitative and nuanced. On the relatively “easy” to define end of the specialization spectrum are obligate biotrophs which must live within plants and therefore have their whole life cycles closely associated with hosts[20]. It is unclear what fraction of the phytobiome exists as biotrophs but many bacteria found within the phytobiome survive quite well *sans* host[16]. This discussion is further compounded by the idea that,to truly define host range, one needs to test many different plants under the same inoculation conditions and that inoculation of plants under laboratory conditions can lead to arguments about what constitutes what constitutes growth or benefits to the host. While there are numerous reports of plant growth promoting bacteria working to provide a benefit on specific host species [21,22], such results might not inherently constitute specialization because the potential for growth on alternative hosts can’t be excluded. Taking this a step further, to date there are few reports that describe the molecular basis for fine scale host association for beneficial endophytic strains outside of nodulating bacteria, although a few higher level trends have been noted [19,23,24]. However, unpacking our current overall understanding does provide a good snapshot for what mechanisms one expects to be involved in establishing interactions: chemotaxis towards specific plant factors, bacterial secretion systems, plant peptide signals, differential protease activity, and general abilities of the symbiont to overcome/avoid plant defenses [25–29].

## Can Plants Shape Their Own Microbiomes?

Once individuals have laid down roots in an area, most plants lack the ability to modify their microbiomes by movement to new areas and are therefore much more dependent than animal microbiomes on being able to recruit from immediate environmental surroundings. However, plants are well known to be able to release root exudates to the surrounding soil, which can function as both a chemoattractant and potentially a prebiotic nutritional source for scavenging microbes[30–32]. That these exudates differ differ between plant species, and indeed within the same species under different conditions [33–35], paints the intriguing narrative that evolution could favor the acquisition of exudate catabolic operons by beneficial bacteria. One particular challenge with the production of exudates to recruit microbes, however, is that it is difficult to police the catabolism of such public goods outside of the host (Box 2).

### Box 2

If plants produce public goods (i.e. root exudates) to attract beneficial environmental bacteria, it is well established in other systems that “cheater” strains will arise to disrupt the partnership. These cheaters can catabolize substrates but don’t pay energetic costs associated with providing benefits, a situation known as the “free rider problem” in social evolution studies. One way for plants to police the evolution of cheaters is to limit the production of public goods so that they can only be utilized by beneficial partners. Since it is easier for plant hosts to police bacterial strains that they directly interact with, reciprocal benefits and coevolution is more likely to occur for endophytic bacterial partners. Indeed, there is growing evidence that plants can sanction which nodules receive benefits depending on the quality of the bacterial symbionts.

Inevitably, microbial symbionts are required to navigate plant immune responses, and in many cases it appears as though the same general defense pathways are involved regardless of whether host plants are dealing with symbionts or pathogens [28]. Given this overlap, there exists potential for specialization and for refining interactions between partners at each step of recognition and response. Plant PAMP/ MAMP receptors provide an initial layer of surveillance, and establish the baseline abilities for microbes to interact closely [36]. Since each plant species can survey for numerous molecular signatures of microbes, and there is diversity across plant species for which epitopes (and even flavors of epitopes for the same protein) are recognized, it likely that that phytobiome interaction landscapes differ between plants in evolutionarily relevant ways [28,37,38]. It is unclear how many potential beneficial symbionts are recognized during this first line of defense, and what such recognition means for any downstream beneficial effects, but it is notable that some nodule forming symbionts can avoid MAMP recognition pathways[38]. There is also precedent for non-pathogenic microbes to be limited in growth by downstream defense responses such as through R-gene mediated defense[25]. If R-genes and the hypersensitive broadly affects members of the phytobiome, it will be interesting to see how differences in the activity of these type of defenses between plant tissue compartments could shape microbiome membership. Lastly, given the production of plant hormones and hormone mimics by phytobiome members and that tradeoffs between plant immune responses and growth are mediated by complex signalling pathways, great potential exists for phytobiome communities to regulate and be regulated by complex feedback loops which integrate plant stress signals[39,40].

There was, and remains, a hope that plant breeding technologies can be used to identify genes that control phytobiome development and which could hold the keys to community engineering. However, somewhat blunting these hopes, is the widespread result that genotype by environment interactions appear to be rampant for these phytobiome associated traits[13,41]. A clear demonstration of this effect can be seen in the intimate interaction between Ensifer and nodulation of Trifolium, as fertilizer use is associated with decrease in nodulation [42]. Moreover, such changes could create cascading effects as nodulation status appears to shape other broad interactions across the microbiome [43]. Cascading effects are not restricted to nodulation as certain microbial species appear to also act as “hubs” to shape and pattern the phytobiome in Arabidopsis. Similar effects can be seen in maize using simplified communities [44]. Highlighting the challenges of trying to map and associate plant genetic determinants with microbiome composition, a large-scale study by in maize demonstrated that, while there were genetic components that solidly determined microbiome composition across fields, genetic background had very little explanatory power for the microbiome as a whole[13].

## Coevolution Within Microbiomes

While likely a product of ease of language, there is a tendency across microbiome literature to attribute correlations in presence of microbes on a particular host species to coevolution. Following a strict point of view, coevolution requires one to demonstrate a precise series of genetic changes in host and symbiont that directly occur in response to one another[45]. A looser definition can be employed, which takes into account correlations in genetic backgrounds through space and time without definitively proving evolutionary linkage[46]. Somewhat looser still, and approached more from a phenomenological standpoint, work on animal microbiomes have shown that microbial communities can co-diverge with their hosts in a process termed phylosymbiosis[47]. A challenge when discussing both of these latter situations is that, although both *could* describe a coevolutionary process, other evolutionary dynamics could give rise to similar population patterns.

It’s possible that coevolutionary scenarios qualitatively differ between beneficial symbionts and those of pathogens. When discussing pathogens, oftentimes the “Red Queen” hypothesis is invoked whereby there is an arms race between host and pathogen due to the evolution of new levels of immune response and countermeasures by the bacteria[48]. An arms race isn’t expected to occur for beneficial microbes because the immune system is evolving to work with the symbiont rather than defend against it, but a “Red King” is likely more relevant for beneficial microbes in phytobiomes[49]. Under this scenario, microbes would often be left waiting for the plant host to adapt because the slowest partner controls the evolutionary dynamics. Such a situation might bias the potential for coevolution between phytobiome and plant host to those species that have shortest generation times rather than longer lived plants since their evolutionary response rates are slightly closer to their associated microbes.

Since microbes are going to be growing and thriving in the context of other phytobiome members,coevolutionary scenarios may be more relevant to understanding how other members of the phytobiome respond to each other rather than to the host plant itself. Indeed, there is a growing body of evidence that suggests that bacteria found within plants are coevolving in response to each other[50], with fungi[51], as well as to phage present within these populations[52]. There have been a variety of molecular mechanisms discovered whereby bacteria can specifically target other microbes present within the phytobiome to kill or aid[53–55], and such intra and interspecies interactions could be a major force structuring microbiome communities. Even at this level, demographic patterns could strongly shape the ability of microbes to coevolve with each other. One might expect bacteria with correlated patterns of transmission across plant hosts to interact more strongly over evolutionary time than microbes that only occasionally encounter each other. Furthermore, while annuals grow and reproduce during a single season and thus are likely affected by whatever microbes happen to colonize during that single bout of growth, perennials spring up year after year in the same area. Given these demographic differences, perennial hosts might be able to better cultivate and nurture their microbial surroundings. It is also a strong possibility that root/bulb endophytes play an outsized role by acting as “refugia” for perennial species in between growing seasons.

## Conclusions

Phytobiomes shape plant developmental and evolutionary dynamics, and thus a deeper understanding of the forces that structure plant-associated microbes could enable engineering of these communities to boost agricultural yields. It is now clear that all plants harbor a vast diversity of microbes throughout all stages of their life cycle, and that microbes are continually entering and exiting from these communities. While there exists evidence that specialization can occur whereby microbes only colonize specific hosts and by which plants can influence their microbes, these relationships appear to be most heavily influenced by environmental factors. Likewise, it remains a possibility that coevolution has occurred between plants and some of the microbiome members, but proving such dynamics will be difficult given that many phytobiome members can survive and thrive outside of host plants and often on a variety of hosts. Overall, we are likely closing in on the end of the beginning of phytobiome research, whereby there is solid foundation of knowledge to predict for what kinds microbes will be found across hosts under different conditions, but many questions concerning evolutionary dynamics of phytobiomes remain to be answered in the years to come.

**Figure 1.**
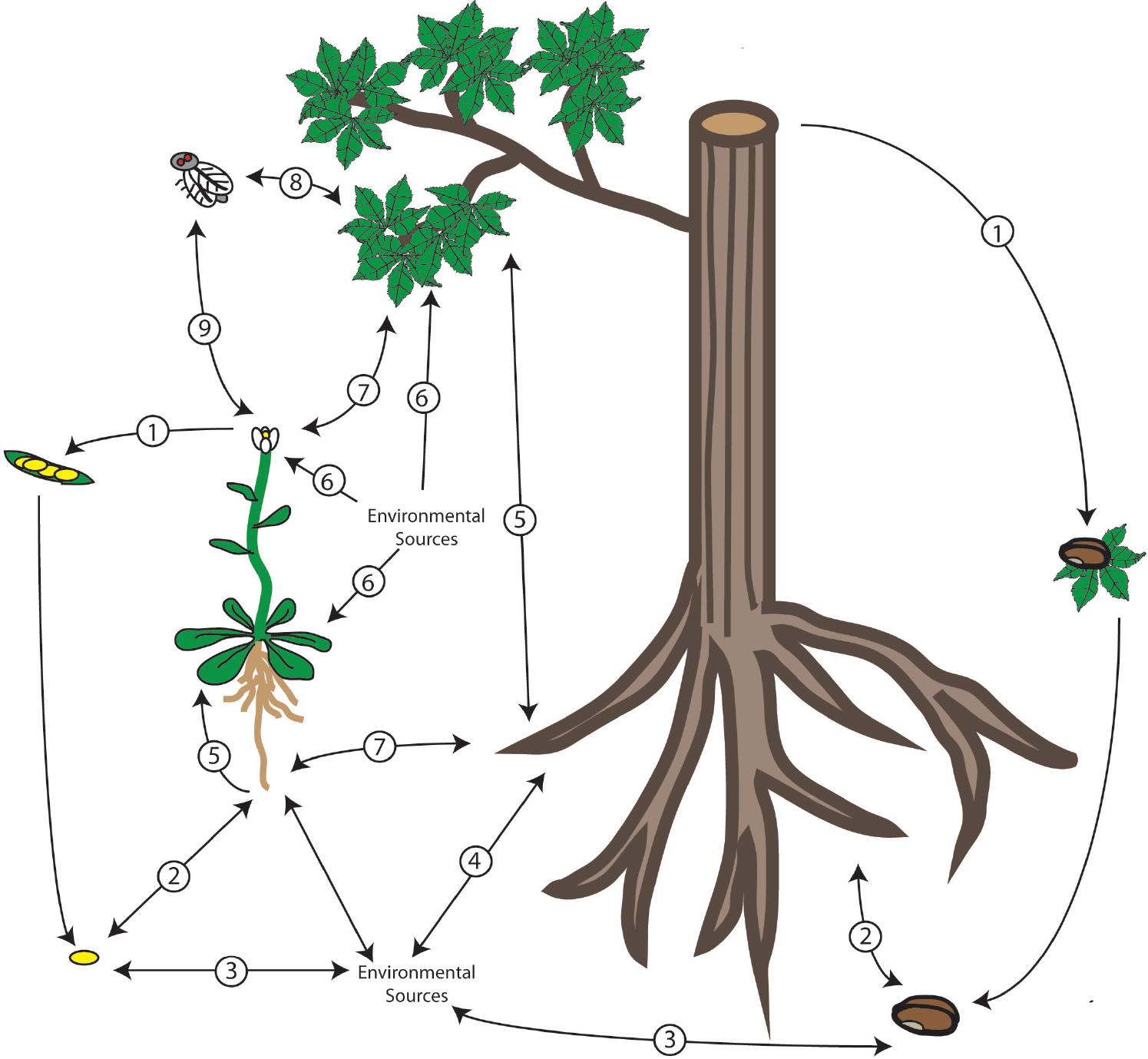
Phytobiome Recruitment and Establishment is a Complex Process. There are numerous inputs for new microbial strains and species to join phytobiome communities. The influence of each of these routes will likely change over the course of an individual plant’s life cycle. Various demographic and developmental factors could also greatly skew phytobiome recruitment and succession patterns such as whether plants are annuals and perennials. Numbered routes in the figure represent a subset of these possibilities and include: 1) vertical transmission from plant to seed 2) invasion of seed endosphere from root endophytes and vice versa 3) invasion of seed endosphere from environmental sources 4) invasion of roots from environmental sources 5) leave colonization from root associated microbes 6) leaf and flower colonization from environmental sources 7) cross-species colonization 8) insect vectoring 9) movement of pollen microbiomes by pollinators

**Figure 2.**
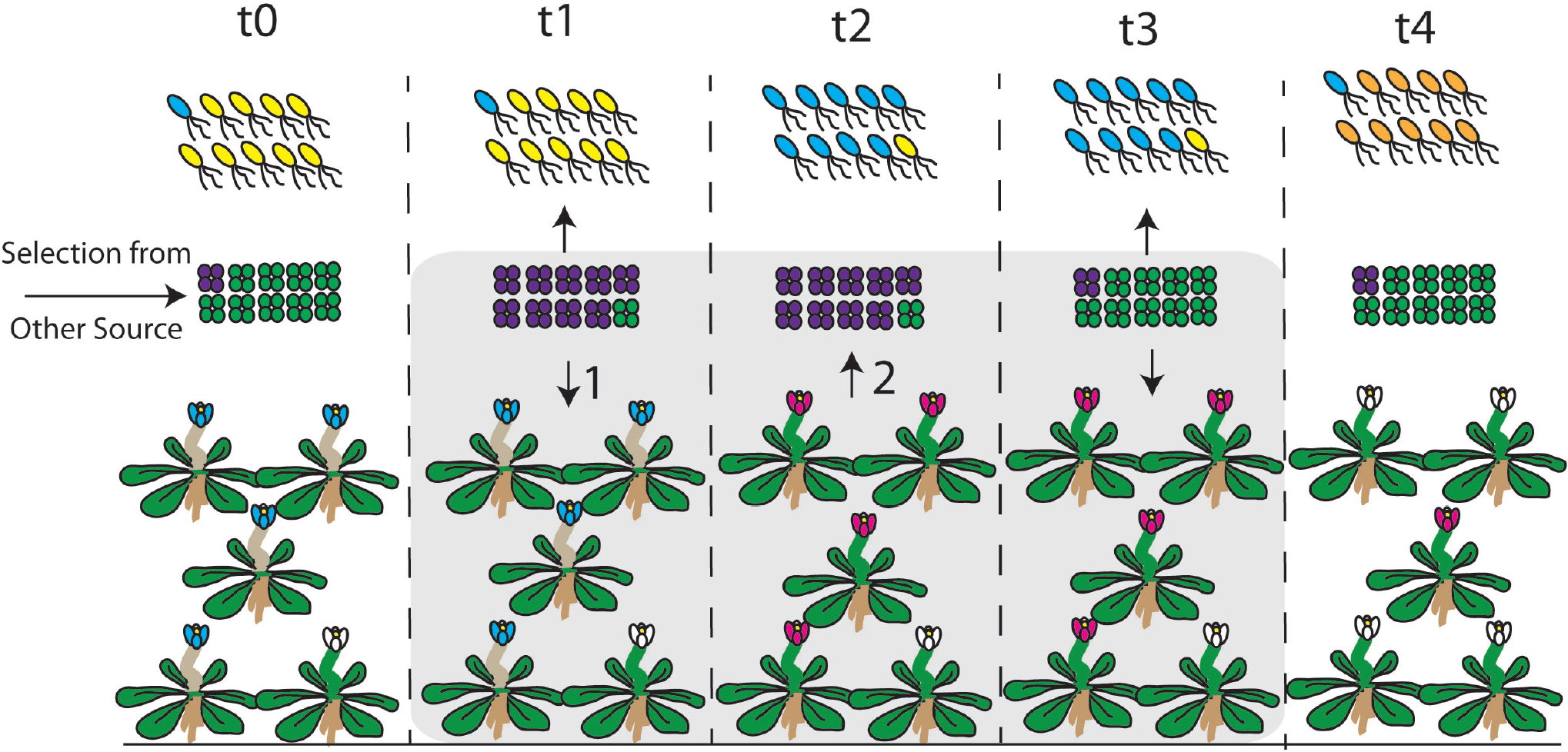
The Challenge of Interpreting Coevolutionary Patterns. Although numerous studies refer to coevolutionary relationships between hosts and microbes, interpretation of population-wide patterns in a coevolutionary context can be difficult. Here I illustrate sampling from 5 different time points of the same phytobiome and host populations to highlight when it is clear that coevolution has occurred but also to demonstrate how correlational patterns can confound analyses. Three different organismal populations that make up a phytobiome (rod on top, cocci in middle, flower on bottom) are represented with arrows denoting selective forces. Coevolution is strictly defined as the reciprocal evolution of at least two organisms in response to one another, and in this figure, three time points are minimally required to properly interpret coevolution. At t1, flower genotype is directly influenced by cocci genotype (arrow #1), while at t2 flower genotype reciprocally influences cocci genotype (arrow#2). Finally, the reciprocal evolutionary response of the cocci genotype is witnessed at t3. Interpretation of this pattern is confounded, however, because other selective forces influence cocci genotype between t0 and t1. Furthermore, even though the rod genotype on top is solely influenced by selection from the cocci strains even though genotype frequencies correlate exactly with those of the host plant.

**Figure 3.**
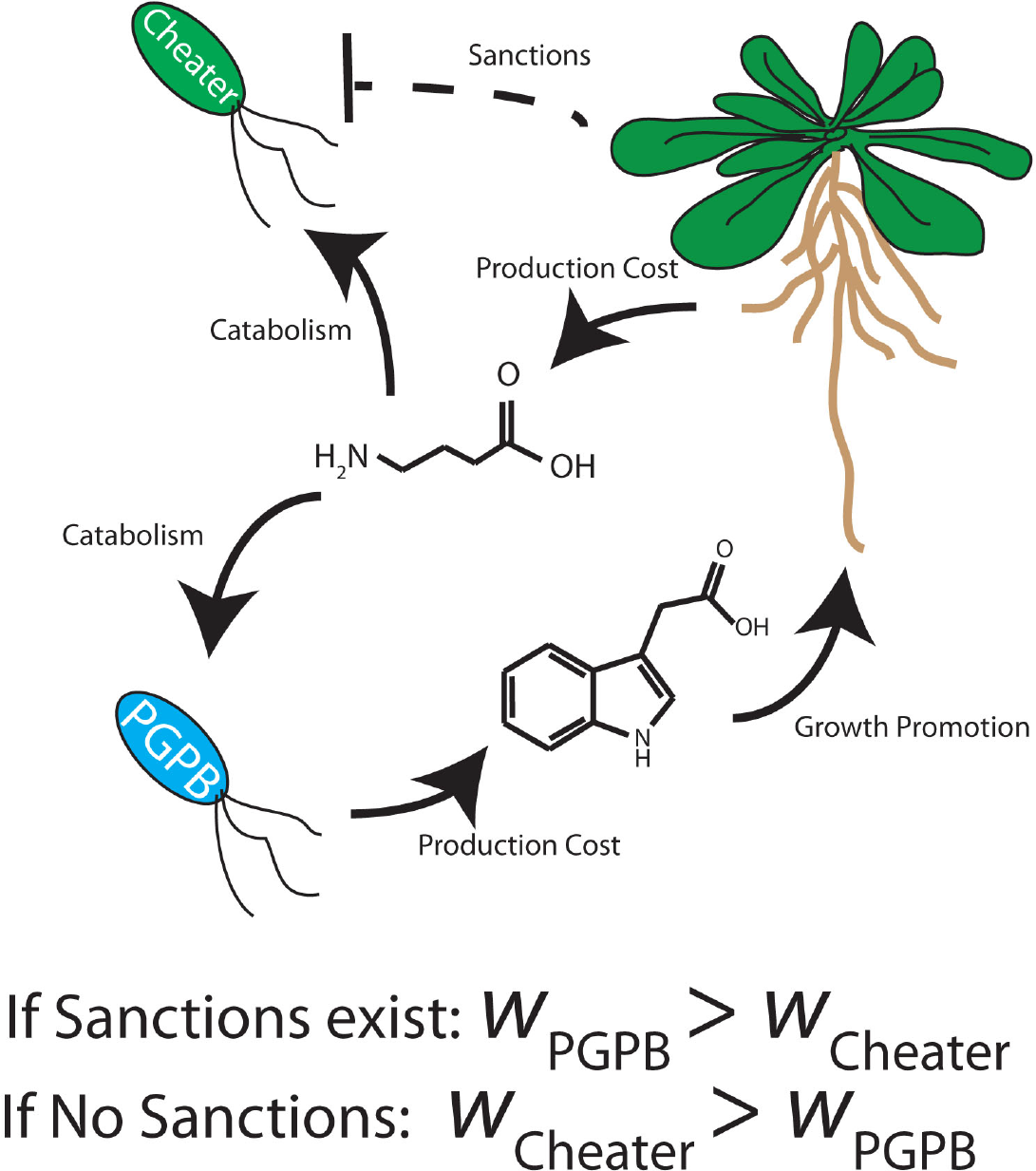
The Free Rider Problem with Root Exudates. If there publicly available goods, in this example GABA as a root exudate, are produced to lure in beneficial microbes like PGPB that produce IAA, the system could breakdown due to the evolution of cheater cells. Here, cheater cells can catabolize GABA but do not pay an energetic cost of having to produce IAA and thus can exploit the plant host. Sanctioning the use of the public good so that GABA can only be catabolized by IAA producing partners provides a means for plants to police the evolution of cheater cells.

